# A role for BCL2L13 and autophagy in germline purifying selection of mtDNA

**DOI:** 10.1101/2022.09.02.506367

**Authors:** Laura S. Kremer, Lyuba V. Bozhilova, Diana Rubalcava-Gracia, Roberta Filograna, Mamta Upadhyay, Camilla Koolmeister, Patrick F. Chinnery, Nils-Göran Larsson

**Affiliations:** Department of Medical Biochemistry and Biophysics, Karolinska Institutet, Stockholm, Sweden; MRC Mitochondrial Biology Unit, University of Cambridge, Cambridge, United Kingdom; Department of Clinical Neuroscience, School of Clinical Medicine, University of Cambridge, Cambridge, United Kingdom

## Abstract

Mammalian mitochondrial DNA (mtDNA) is inherited uniparentally through the female germline without undergoing recombination. This poses a major problem as deleterious mtDNA mutations must be eliminated to avoid a mutational meltdown over generations. At least two mechanisms that can decrease the mutation load during maternal transmission are operational: a stochastic bottleneck for mtDNA transmission from mother to child, and a directed purifying selection against transmission of deleterious mtDNA mutations. However, the molecular mechanisms controlling these processes remain unknown. In this study, we systematically tested whether decreased autophagy contributes to purifying selection by crossing the C5024T mouse model harbouring a single pathogenic heteroplasmic mutation in the tRNA^Ala^ gene of the mtDNA with different autophagy-deficient mouse models, including knockouts of *Parkin, Bcl2l13, Ulk1*, and *Ulk2*. Our study reveals a robust effect of knockout of *Bcl2l13* on the selection process, and weaker evidence for the effect of *Ulk1* and potentially *Ulk2*, while no statistically significant impact is seen for knockout of *Parkin*. This points at distinctive roles of these players in germline purifying selection. Overall, our approach provides a framework for investigating the roles of other important factors involved in the enigmatic process of purifying selection and guides further investigations for the role of BCL2L13 in the elimination of non-synonymous mutations in protein-coding genes.

## Introduction

Mutations in mtDNA are an important cause of mitochondrial disorders where about one in 200 healthy individuals carries a pathogenic mutation in mtDNA and one in 10000 individuals is affected by mtDNA disease (1). Mammalian mtDNA is a circular genome of about 16.6 kb size can exist in hundreds to thousands of copies per cell. If all mtDNA copies have the same genotype the state is called homoplasmy, whereas heteroplasmy indicates the co-existence of different genotypes in the same cell, e.g., wild-type and mutant mtDNA. The mtDNA is inherited uniparentally through the female germline (2) without undergoing recombination (3). This poses a major problem, as deleterious mtDNA mutations must be eliminated to avoid a mutational meltdown over generations (4). Two processes that operate to decrease the mutation load during maternal transmission have been identified, i.e., (i) the genetic bottleneck, which results in the stochastic transmission of only a subset of the pool of mtDNAs present in the mother, and (ii) purifying selection, which prevents transmission of deleterious mtDNA mutations. The mechanisms regulating these processes are unknown, and it is also unclear whether the bottleneck and purifying selection processes are linked to each other.

A possible link between the genetic bottleneck and purifying selection is supported by recent evidence from the C5024T mouse model harbouring a pathogenic C to T substitution within the mitochondrial tRNA^Ala^ gene. In this mouse model, mean heteroplasmy levels are consistent across all tissues (5,6), which simplifies comparisons of heteroplasmy levels between mothers and offspring. When breeding C5024T females, no pups with a heteroplasmy level higher than 80% are obtained (7). However, increasing mtDNA copy number in the mother via TFAM overexpression shifted the tolerated mutation level in the offspring to 84% (5). This finding shows that the mtDNA copy number influences purifying selection of mtDNA. The actual mechanism for purifying selection is not yet known, but likely involves a functional test of mtDNA and its ability to sustain normal oxidative phosphorylation (OXPHOS) at the organellar, or cellular level (8). One appealing hypothesis is that mtDNA mutations impair OXPHOS, which, in turn, leads to depolarization of the mitochondrial inner membrane due to the inability to sustain a normal membrane potential. This decline in OXPHOS function may facilitate subsequent removal of the depolarized mitochondrion by autophagy (also known as mitophagy).

In mammals, several pathways are in place for the removal of mitochondria via autophagy, these might be partially redundant to ensure robustness of the process or allow the removal of mitochondria under specific conditions (9–11). There are two main pathways that specifically can target mitochondria for degradation by the lysosome involving either ubiquitin- or adapter-dependent marks on the outer mitochondrial membrane. PARKIN plays an important role in the former pathway as it functions as a E3 ubiquitin ligase (12). Knockout of *Parkin* in mice harboring random mtDNA mutations has been suggested to affect the pathogenicity of mtDNA mutations in the striatum and it could possibly also play a role in controlling the inheritance of mtDNA (13). A representative of the adapter-dependent pathway is BCL2L13, the mammalian homologue of ATG32, which is essential for the autophagic removal of mitochondria in budding yeast (14–16). Intriguingly, *Bcl2l13* expression can rescue the mitophagy phenotype caused by *Atg32* knockout in yeast (14). Eventually, most of the mitophagy pathways converge when the mitochondrion is delivered to be engulfed by the autophagosome. Autophagosome formation is initiated by the serine/threonine protein kinase ULK1, the mammalian homologue of ATG1 (17–19). It was recently reported that knockdown of the *Drosophila melanogaster* homologue of *Ulk1* is necessary for germline selection of mtDNA in the fly (20). Importantly, ULK2, an ULK1 homologue, shares about 52% protein sequence identity with ULK1 and it is thought that these two proteins act redundantly in some tissues, while in other tissues they might have non-overlapping roles (18,21). In the current study, we investigated the role of PARKIN, BCL2L13, ULK1, and ULK2 in transmission and purifying selection in the C5024T mouse model.

## Results

### Gene expression levels in the germline

For PARKIN, BCL2L13, ULK1, or ULK2 to contribute to purifying selection, they need to be expressed when selection occurs. Segregation of mtDNA likely occurs during germ cell development, the exact timing is however not known. We therefore checked the expression of the genes encoding these autophagy factors at various stages during germ cell development in publicly available databases and datasets. As these proteins are conserved in mouse and humans, we queried mouse as well as human data. According to the GTEx Portal (GTEx Analysis Release V8 (dbGaP Accession phs000424.v8.p2)) (22) and the Human Protein Atlas (23), PARKIN, BCL2L13, ULK1, and ULK2 are expressed in the adult human ovary. Furthermore, expression of *Bcl2l13, Ulk1*, and *Ulk2* transcripts was detected in mouse E12.5 primordial germ cells (PGCs), female germline stem cell (FGSCs), germinal vesicle (GV) oocytes and metaphase II (MII) oocytes, whereas expression of *Parkin* (also known as *Park2)*, the gene encoding PARKIN, was detected in all stages besides PGCs (24). As the genes seemed to be readily expressed at various stages of germ cell development in the female germline, they could play a role in the purifying selection of mtDNA.

### Knockout of *Parkin, Bcl2l13, Ulk1*, or *Ulk2* in C5024T females does not shift the maximal tolerated mutation level in their offspring

If the removal of mitochondria via autophagy contributes to mtDNA selection, we would expect that knockout of key players in the process would result in a weaker selection. Weaker mtDNA selection, in turn, can manifest in different ways. One possible effect of weaker selection is an increase in the maximal tolerated levels of mutated mtDNA in heteroplasmic mouse pups. This was previously reported to occur as a result of increasing the mtDNA copy number via TFAM overexpression in the C5024T mice (5). We therefore investigated whether knockout of *Parkin, Bcl2l13, Ulk1*, or *Ulk2* shifted the maximally tolerated mutation load in the offspring of heteroplasmic C5024T mice. To this end, we generated whole body *Parkin*^-/-^ C5024T, *Bcl2l13*^-/-^ C5024T, *Ulk1*^-/-^ C5024T, and *Ulk2*^-/-^ C5024T females along with *Parkin*^*+/+*^ C5024T and *Bcl2l13*^*+/+*^ C5024T litter mate control females, all on the same mtDNA and nuclear C57BL/6NCrl genetic background (Fig. 1A). As the whole body *Bcl2l13* knockout mouse model was newly generated by us, we confirmed that the protein was indeed absent in livers isolated from females after mating (Fig. 1B). The *Bcl2l13* knockout mouse model is viable, fertile, and does not develop any obvious phenotype. For each group, we included at least 5 females, all of which had a heteroplasmy level of more than 55% where purifying selection reportedly occurs on transmission (7). We subsequently crossed these females to C57BL/6NCrl males (Fig. 1A) and analyzed the mutation load in the resulting pups.

**Figure 1.**
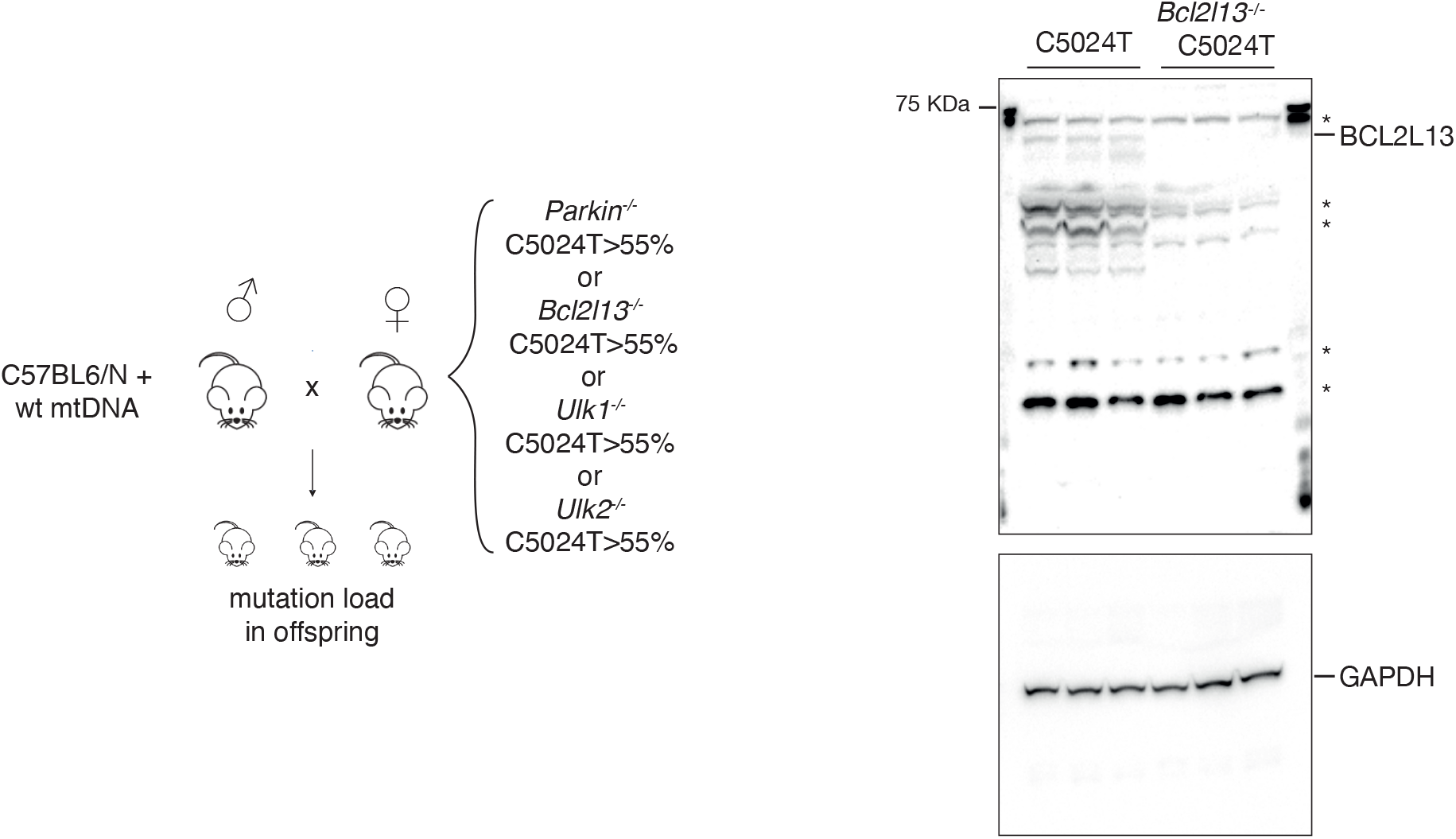
**A** Overview of the experimental setup. Female mice transmitting the pathogenic C5024T mutation in combination with Parkin^-/-^, Bcl2l13^-/-^, Ulk1^-/-^, or Ulk2^-/-^alleles were crossed to C57BL6/N males with wild-type (wt) mtDNA and the mutation load was measured in the resulting offspring. **B** Western blots analysis of BCL2L13 protein levels in mouse liver (n=3; *= unspecific bands), levels of GAPDH shown as loading control.

In total, we measured the heteroplasmy level from 959 pups born to 40 mothers (Table 1). As we did not observe any difference in heteroplasmy distribution between the *Parkin*^*+/+*^ C5024T and *Bcl2l13*^*+/+*^ C5024T control groups (Kolmogorov–Smirnov test p-value = 0.31) and wanted to keep the animal use to a minimum, we did not generate *Ulk1*^*+/+*^ C5024T or *Ulk2*^*+/+*^ C5024T control groups. Instead, throughout our analysis we used both the *Bcl2l13*^*+/+*^ C5024T and *Parkin*^*+/+*^ C5024T cohorts as controls. It must be noted that out of all samples measured, one sample belonging to the *Bcl2l13*^*+/+*^ C5024T group exceeded the 80% threshold. This turned out to be a technical artefact as remeasurement of the sample revealed a heteroplasmy level below the threshold (1035:417 pup heteroplasmy (%): original measurement: 86; repeated measurement: 71). We did however not want to manually remove this outlier and it is hence included in the analysis. Analysis of the maximally tolerated mutation load in the different groups revealed that neither knockout of *Parkin, Bcl2l13, Ulk1*, nor *Ulk2* in C5024T mothers had any effect on the maximal tolerated heteroplasmy (Figure 2). This was in contrast to what was observed previously when increasing the mtDNA copy number employing a similar number of breeding pairs and pups (5).

**Table 1.** Mothers and pups in each genotype group. Pup heteroplasmy and mother-to-pup normalised heteroplasmy shift are given as mean ± st.dev.

| Genotype group | # mothers | # pups | Pup heteroplasmy (%) | Normalised heteroplasmy shift |
| --- | --- | --- | --- | --- |
| **Parkin:+/+** | 4 | 122 | 69.73±6.04 | -0.20±0.29 |
| **Bcl2l13:+/+** | 6 | 152 | 68.70±6.61 | -0.12±0.33 |
| **Parkin:-/-** | 5 | 159 | 69.40±6.57 | -0.20±0.34 |
| **Bcl2l13:-/-** | 9 | 236 | 69.19±5.76 | -0.01±0.32 |
| **Ulk1:-/-** | 10 | 148 | 69.93±6.16 | -0.06±0.31 |
| **Ulk2:-/-** | 6 | 142 | 68.84±7.11 | -0.08±0.33 |

**Figure 2.**
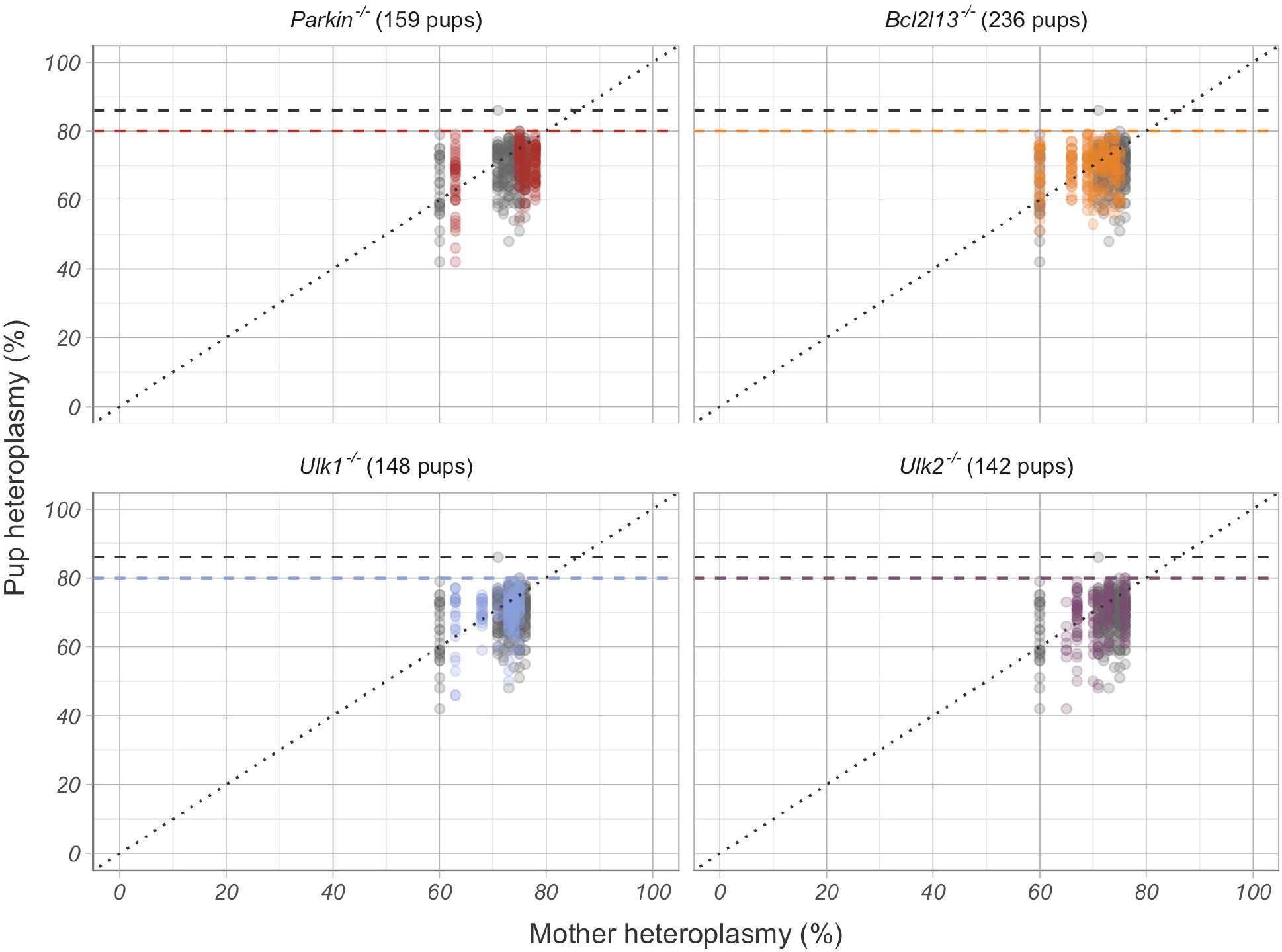
Mother and pup C5024T heteroplasmy across Parkin, Bcl2l13, Ulk1 and Ulk2 knockout groups. Coloured points correspond to mice with autophagy-mediating gene deletion and grey points correspond to mice from both the Parkin^+/+^ and Bcl2l13^+/+^ controls (274 pups in total). The dashed lines show the highest observed heteroplasmy level in each group (80% for Parkin, Ulk1 and Ulk2 knockout groups, 79% for the Bcl2l13 knockout group, and the outlier 86% for Bcl2l13^+/+^). The highest observed heteroplasmy was comparable across all groups.

### Knockout of *Bcl2l13, Ulk1*, and *Ulk2* affect the heteroplasmy shift from mother to offspring in C5024T mice

Weaker mtDNA selection can skew the overall pup heteroplasmy distribution towards higher values, without necessarily changing the maximal heteroplasmy. Pup heteroplasmy is not independent of maternal heteroplasmy (25), and therefore differences in maternal heteroplasmy across groups may confound any comparative analysis. As it is not possible to strategically breed mothers with the exact same heteroplasmy level for the different groups (Fig. 3A), we instead calculated a mother-to-offspring normalized heteroplasmy shift from the observed heteroplasmy measurements which accounts for this confounder in the analysis (Fig. 3B–C) (6,25). In the absence of selection pressure, the mean heteroplasmy shift is zero (26), but if the mean heteroplasmy shift is negative, this is indicative of purifying selection. In agreement with previously reported results (25), purifying selection was present in high-heteroplasmy C5024T mothers across both *Parkin*^*+/+*^ C5024T and *Bcl2l13*^*+/+*^ C5024T control groups, where the mean heteroplasmy shift was -0.20 ± 0.29 and - 0.12 ± 0.33 (mean ± st.dev.) respectively (Table 1, Fig. 3C). A similar negative shift was also observed in the *Parkin* knockout group (−0.20 ± 0.34). However, the shift in the *Bcl2l13* knockout group was closer to zero (−0.01 ± 0.32), and shifts in the *Ulk1* and *Ulk2* knockouts were higher than in both controls (−0.06 ± 0.31 and -0.08 ± 0.33, respectively).

**Figure 3.**
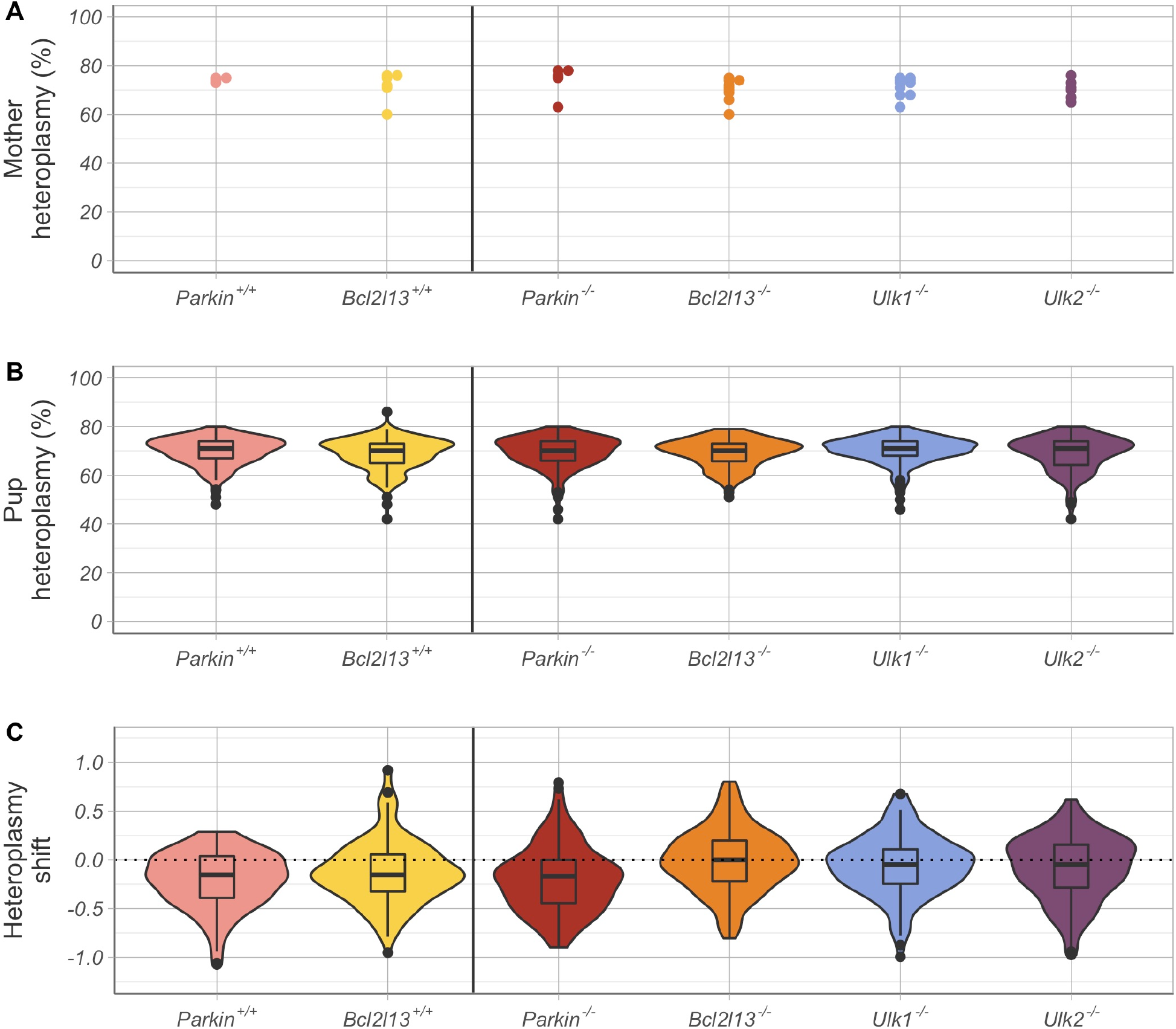
Mother and pup heteroplasmy across the six groups. **A**. High-heteroplasmy (60–78%) mothers were selected in each group. **B**. Pups within each group exhibited comparable high heteroplasmy (69.3 ± 6.3% across the cohort). **C**. The mean normalised heteroplasmy shift was negative across all groups. The shift was closest to zero for the Bcl2l13 (−0.01 ± 0.32; orange) and Ulk1 (−0.06 ± 0.31; blue) knockout groups, indicating that selection may be partially disrupted by the absence of functional BCL2L13 and ULK1.

This initial exploratory analysis suggests that knockouts of *Bcl2l13, Ulk1* and *Ulk2* have effect on the mean heteroplasmy shift, and therefore on purifying selection. Wilcoxon’s rank tests, with Bonferroni multiple-testing correction show that the shifts in the *Parkin*^*+/+*^, *Bcl2l13*^+/+^, and *Parkin*^-/-^ groups were significantly different from zero (p-value < 0.001 in all cases), while the shifts in *Bcl2l13*^-/-^, *Ulk1*^-/-^, and *Ulk2*^-/-^ were not (p-values 0.678, 0.040, and 0.015 respectively). However, in the data, mothers have multiple pups (23.89 *±* 10.64), and therefore each mother-offspring observation is not independent, violating standard statistical hypothesis test assumptions. For example, any measurement errors on the maternal heteroplasmy levels could propagate across tens of observations and bias the results.

In order to robustly test whether the effect of knockout of *Bcl213, Ulk1*, and *Ulk2* on the normalized heteroplasmy shift was statistically significant, we performed Kolmogorov–Smirnov tests comparing gene knockout groups to subsamples of the control data. Testing the larger of the control groups, *Bcl2l13*^*+/+*^ C5024T (six mothers, 152 pups), against all other groups indicated that only the *Bcl2l13* knockout resulted in significantly different distribution of heteroplasmy shifts (p-value < 0.001). To ascertain that our results were not because we restricted the control data to the six *Bcl2l13*^*+/+*^ C5024T mothers, we further generated all 210 possible subsampled control datasets, consisting of any six mothers and all their respective offspring from the *Bcl2l13*^*+/+*^ C5024T and *Parkin*^*+/+*^ C5024T groups. For example, one such subsampled control could consist of all four mothers in the *Parkin*^*+/+*^ C5024T group and two out of the six mothers in the *Bcl2l13*^+/+^ C5024T group. Another subsampled control could consist of one mother from the *Parkin*^*+/+*^ C5024T group and five mothers from the *Bcl2l13*^*+/+*^ C5024T group. We subsequently performed Kolmogorov–Smirnov tests for the *Parkin*^*+/+*^ C5024T and *Bcl2l13*^*+/+*^ C5024T control groups and each knockout group against each of the 210 subsampled controls. We measured how frequently the genotype groups were significantly different to the subsampled controls (p-value < 0.008 after Bonferroni multiple-testing correction). The proportion of subsampled control tests resulting in p-values below 1% and below 5% for each genotype group are depicted in Figure 4A. As expected, the *Parkin*^*+/+*^ C5024T and *Bcl2l13*^*+/+*^ C5024T control groups did not significantly differ from any of the 210 subsampled controls (see also Figure 4A, left and middle upper panel). Significant differences of the *Parkin* knockout group were rare (14 out of 210 tests, or 6.67% of tests), indicating the knockout is indeed unlikely to affect the heteroplasmy shift (Figure 4A, right upper panel). In contrast, the *Bcl2l13* knockout group was significantly different from the vast majority of subsampled controls (195 out of 210 tests, or 92.86% of all tests), providing robust evidence of the *Bcl2l13* effect on purifying selection (Figure 4A, left lower panel). For the *Ulk1* group, over half of tests showed significant difference (120 out of 210 tests, or 57.14% of all tests), with median p-value = 0.005 below the significance threshold (Figure 4A, middle lower panel). Results for the *Ulk2* group were similar to the results for *Ulk1* but weaker with less than half (90 out of 210 tests, or 42.86% of all tests) showing a statistically significant difference, with a median p-value = 0.013 above the threshold (see also Figure 4A, right lower panel).

**Figure 4.**
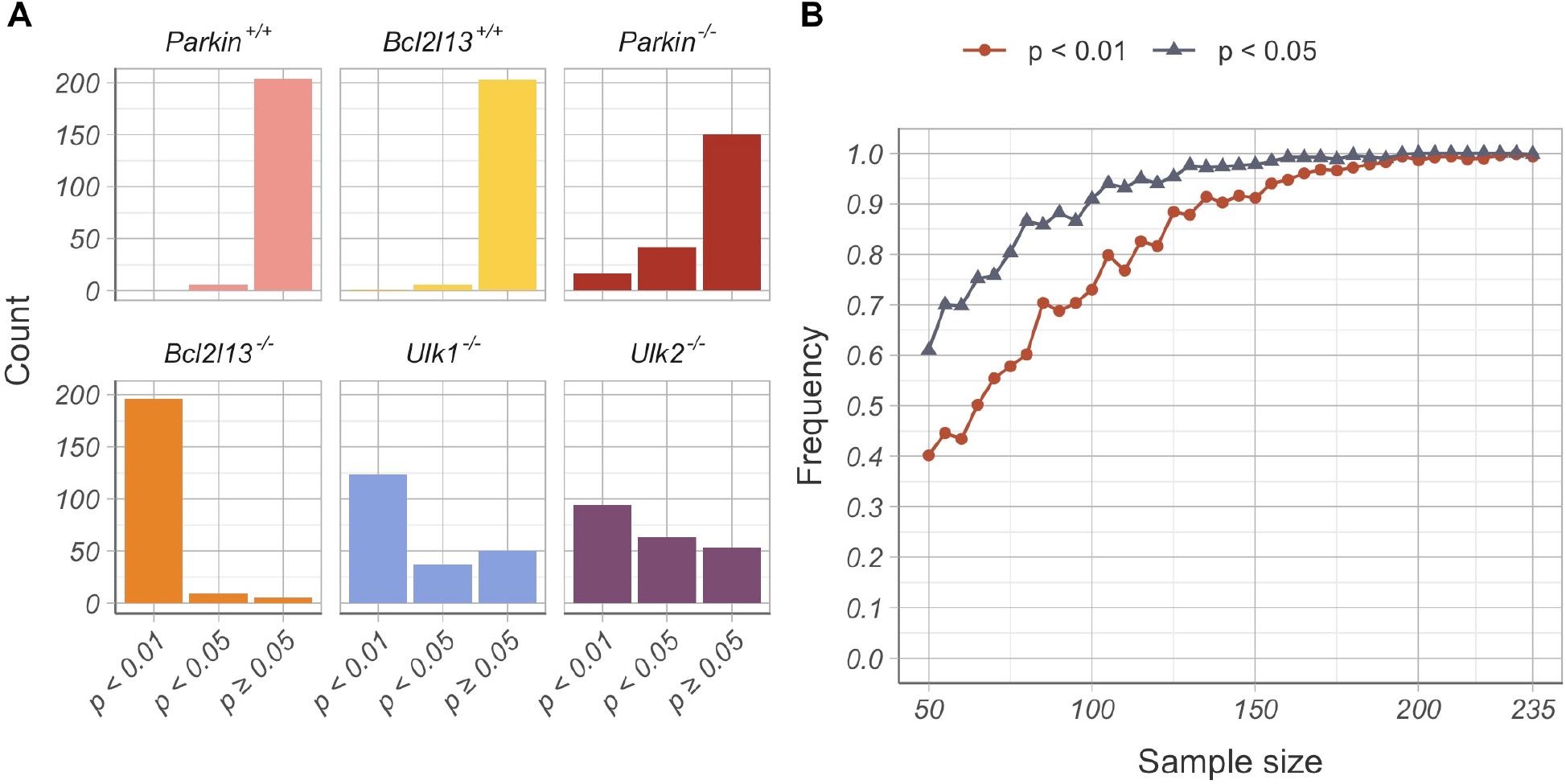
Robustness analysis of results. **A** Proportion of bootstrap p-values showing a significant difference between subsampled control data and the six experimental groups. As expected, Parkin (pink) and Bcl2l13 (yellow) wildtype groups were not significantly different to most subsampled controls. The Parkin knockout (red) also had no significant effect on heteroplasmy shift. The Bcl2l13 knockout group (orange) was consistently significantly different from subsampled controls. Tests for the Ulk1 (blue) and Ulk2 (purple) knockouts showed more variable results, with the effect of Ulk1 knockout being significant in over half of samples. **B** Subsampling of the Bcl2l13 knockout data and controls shows the effect is consistently observed at lower sample sizes.

Finally, we wanted to ascertain whether the effect observed for the knockout of *Bcl2l13* was not simply due to the bigger sample size in this group (236 pups compared to 122–159 pups in other groups). We therefore repeated the Kolmogorov–Smirnov test comparing *Bcl2l13* ^-/-^ C5024T to controls by resampling both knockout and control data with replacement. Knockout and control subsamples of different sizes were repeatedly generated, and the fraction of significant differences for each sample size was calculated (Figure 4B). For sample size n = 120, the effect of *Bcl2l13* knockout is already reliably detected in 81.6% of replicates at the 1% significance level. Therefore, we concluded that for all our knockout groups our sample size was sufficient to detect a statistically significant difference in the distribution of normalized heteroplasmy shift.

In summary, our study reveals a robust effect of knockout of *Bcl2l13* on the selection process, and weaker evidence for an effect of *Ulk1* and potentially also for *Ulk2*, while no statistically significant impact is seen for knockout of *Parkin*.

## Discussion

Inherited mutations of mtDNA are a major cause of rare mitochondrial disorders and are also implicated in the aging process itself as well as in common degenerative diseases and cancer (1,27–29). Importantly, the incomplete understanding of the very different genetics of mtDNA disorders largely prevents a similar progress as has been made in diagnostics and genetic counseling for disease caused by mutations in nuclear genes. A thorough understanding of mtDNA transmission is not only of utmost importance to provide families at risk of transmitting mtDNA disorders with advice about the recurrence risks, but understanding the mechanisms may also provide clues to prevent disease recurrence and limit the impact of mtDNA mutations on common disorders.

Although the first steps towards manipulating heteroplasmy transmission have been made by experimentally deciphering the role of increasing mtDNA copy number on the propagation of the C5024T mutation (5), modulating the mtDNA copy number in human germ cells is not currently possible, making it important to explore other strategies. One of these approaches could be to manipulate bulk or selective autophagy of mitochondria. To explore the connection between autophagy and mtDNA purification, we systematically diminished autophagy at various levels in the C5024T mouse model and performed a comprehensive analysis of the possible effects on purifying selection. Crucially, our analysis takes into account the relatively low number of mothers in the data, and the fact that mother-to-offspring observations are not independent. Our study reveals a robust effect of knockout of *Bcl2l13* on the selection process, and weaker evidence for an effect of *Ulk1* and potentially also for *Ulk2*, while no statistically significant effect was seen for knockout of *Parkin*. This indicates distinctive roles of the different players in purifying selection. BCL2L13 can rescue knockout of *Atg32* and restore mitophagy in budding yeast (14), suggesting that it has a role for mitophagy in mammals. However, the precise function of BCL2L13 is still unknown and further studies are needed to determine whether enhancing BCL2L13 function can counteract transmission of mutant mtDNA.

Although our study is an important step forward in understanding purifying selection of mtDNA in the female mammalian germline, it has several limitations. First, we only investigated one mutation (C5024T), which is relatively well tolerated in the mouse. The impact of knocking out key autophagy proteins could be different for other mutations, especially for other mutations leading to more severe phenotypes. Secondly, as mentioned above, several pathways exist for the removal of mitochondria via autophagy. Unexplored pathways, mitigated by other players in the adapter-dependent pathway for example, may be fruitful avenues of further investigation (10). Thirdly, for technical reasons we began our exploration by knocking out key autophagy proteins in order to show ‘proof or principle’ that mitophagy is important in heteroplasmy transmission, It will therefore be important to explore ways of enhancing autophagy to see if this suppresses the transmission of mutant mtDNA, Finally, functional redundancy may mean that the simultaneous knockout of several proteins will be required to sufficiently disrupt mitophagy to modulate heteroplasmy transmission. In keeping with this, *Ulk1* knockout and *Ulk2* knockout mice appear to be healthy, but mice with a double knockout of *Ulk1* and *Ulk2* die shortly after birth, showing that at least one of these pathways must be present to allow postnatal development (30). Functional redundancy between ULK1 and ULK2 may also explain why individually they exhibit weaker effects on heteroplasmy transmission. Further experiments are therefore needed to better elucidate the role of autophagy as a contributor to effective selection of mtDNA in the female germline. Moreover, the selective removal of dysfunctional mitochondria is just one of several possible mechanisms governing selection. Other possibilities include positive selection of healthy mitochondria, and mechanisms acting on the cellular level rather than on the organelle level (8,31). These mechanisms are not mutually exclusive, making it possible that several act at multiple levels to ensure faithful propagation of functional mtDNA to the next generation. This would make evolutionary sense given the potentially catastrophic effects of the relentless accumulation of mtDNA mutations down the female germ line, leading to ‘mutational meltdown’ and extinction of the species (Mullers ratchet) (32).

Overall, our findings establish a robust role for BCL2L13 in purifying selection of mtDNA in the female germline, and provide both an entry point to determine the effects of other autophagy modulators, and a methodological approach to further dissect molecular mechanisms for this essential and incompletely understood phenomenon.

## Materials and Methods

### Mice

Whole body *Bcl2l13* knockout mice were generated by the Karolinska Center for Transgene Technologies using the CrispCAS9 system. Two guide RNAs (sgRNA1: TCCTCTACGACTGCGTCTCT, sgRNA2: CTGCAGTCCATGCCAGCGGA) targeting exon 2 and 5 of *Bcl2l13*, respectively, were injected alongside Cas9 protein into C57BL/6NCrl zygotes twice. Out of the 38 animals born, three animals carried deletions around the guide RNA targeting exon 5. Out of these 3 founder animals, the animal carrying a 108 bp deletion (g.34512_34619del, NCBI Gene ID: 94044; g.120847678_120847785del, NCBI Reference Sequence: NC_000072.6) was chosen to establish the *Bcl2l13* knockout mouse line. The C5024T mice were generated previously (7). Whole body *Parkin* knockout mice were obtained by crossing mice with *Parkin* exon 7 flanked by loxP-sites (33) to β-actin-cre transgenic mice (34) followed by backcrossing to C57BL/6NCrl mice (Charles River, Germany). Whole body *Ulk1* and *Ulk2* knockout mice were created by mating *Ulk1*^FL^ and *Ulk*2^FL^ mice (The Jackson Laboratory, stock number 017976 and 017977, respectively) to β-actin-cre transgenic mice (34) and subsequent backcrossing for 5 generations to C57BL/6NCrl mice (Charles River, Germany). Animals were housed in a 12-hours light/dark cycle in standard ventilated cages and fed *ad libitum* with a normal chow diet. The study was approved by the animal welfare ethics committee and performed in accordance with European law.

### Tissue isolation

Animals were euthanized by CO_2_ followed by cervical dislocation. Liver was collected immediately, washed with phosphate-buffered saline (PBS), snap-frozen in liquid nitrogen, and stored at -80°C.

### Protein extractions and immunodetections

Livers were grinded in liquid nitrogen using mortar and pestle. Powder was resuspended in 500 μL of lysis buffer (25mM HEPES, pH 7.5; 5mM MgCl_2_; 0.5mM EDTA; 1x EDTA-free protease inhibitor cocktail (Roche); 1% NP-40 and 140 mM NaCl) and homogenized for 15 sec with the T 10 basic ULTRA-TURRAX® system. Samples were centrifuged at 13000 rpm for 10 min at 4 °C and supernatants were collected. Protein concentration was determined with the BCA protein assay kit (Pierce) and 75 μg of protein was resolved by electrophoresis using 12% Bis-Tris Plus NuPAGE gels (Invitrogen) and transferred to PVDF membranes (iBlot 2 system (Invitrogen)). BCL2L13 was detected (Proteintech 16612-I-ap) and GAPDH (ab8245) was used as a loading control.

### Quantification of the C5024T mutation level

The C5024T mutation level was determined as described previously from DNA isolated from ear clips around the timing of weaning (7). Briefly, a 178-base pair fragment spanning the C5024T site was amplified using the primers 5′Biotin TTCCACCCTAGCTATCATAAGC (forward) and GTAGGTTTAATTCCTGCCAATCT (reverse). PCR products were purified using PyroMark binding buffer (Qiagen) and 1 μl Streptavidin Sepharose TM high-performance beads (GE Healthcare) and denatured with a Pyromark Q24 vacuum workstation (Qiagen). Sequencing was performed using the sequencing primer TGTAGGATGAAGTCTTACA and PyroMark Gold Q24 Reagents (Qiagen) according to the manufacturer’s instructions on a PyroMark Q24 pyrosequencer (Qiagen).

### Statistical analysis

Data analysis and figure generation were performed in R v.4.2.1 using standard libraries.

Mother-to-pup normalized heteroplasmy shift was calculated as

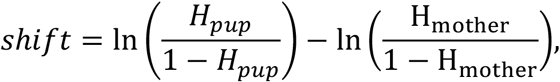

where *H*_*pup*_ and *H*_*mother*_ are the heteroplasmy fractions of the pup and the mother, respectively (24).

Across this study, Wilcoxon’s rank tests and Kolmogorov–Smirnov tests on the heteroplasmy shift were carried out at the 5% significance level. A Bonferroni multiple-testing correction was applied with *n* = 6 when testing all genotype groups simultaneously, and *n* = 5 when testing the *Bcl2l13*^*+/+*^ C5024T control against the other five genotype groups.

When investigating sample size effects, control and *Bcl2l13* knockout data of equal size were generated by sampling 500-fold with replacement. Sample sizes considered ranged from *n* = 50 to *n* = 235 at intervals of five.

## Acknowledgments

We would like to thank Leila Izadi for the support in genotyping of the mice.

## Funding

L.S.K. was supported by an EMBO long-term fellowships (ALTF 570-2019). N.G.L. was supported by grants from the Swedish Research Council (2015-00418), Swedish Cancer Foundation (2021-1409), the Knut and Alice Wallenberg foundation (2016.0050 and 2019.0109), European Research Council (ERC Advanced Grant 2016-741366), Novo Nordisk Foundation, Swedish Diabetes Foundation and grants from the Swedish state under the agreement between the Swedish government and the county councils (SLL2018.0471). PFC is a Wellcome Trust Principal Research Fellow (212219/Z/18/Z), and a UK NIHR Senior Investigator, who receives support from the Medical Research Council Mitochondrial Biology Unit (MC_UU_00028/7), the Medical Research Council (MRC) International Centre for Genomic Medicine in Neuromuscular Disease (MR/S005021/1), the Leverhulme Trust (RPG-2018-408), an MRC research grant (MR/S035699/1), an Alzheimer’s Society Project Grant (AS-PG-18b-022). This research was supported by the NIHR Cambridge Biomedical Research Centre (BRC-1215-20014). The views expressed are those of the author(s) and not necessarily those of the NIHR or the Department of Health and Social Care.

## Author contribution

L.S.K. designed mouse breeding strategies and experiments. L.S.K., D.R.G.M., R.F., and M.U. performed experimental work. L.V.B. and P.F.C. designed the statistical methodology. L.V.B. performed statistical analyses and wrote the data analysis scripts. L.S.K. and L.V.B. analyzed the data, C.K. helped to plan and supervise mouse experiments. N.G.L. conceived and supervised the study. The manuscript was written by L.S.K and L.V.B. All authors read and approved the manuscript.

## Competing interests

N.G.L. is inventor of the C5024T mutant mouse licensed to the pharmaceutical industry by the Max Planck Society. N.G.L. is a scientific founder and holds stock in Pretzel Therapeutics Inc.

## Data and materials availability

All data needed to evaluate and conclude the findings in this paper are present in the paper. All R scripts required to replicate the results and data figures presented can be found at https://github.com/lbozhilova/bcl2l13-mtdna-selection. Requests for the C5024T mouse model or the Bcl2l13 mouse model should be submitted to N.G.L.

## Notes

https://github.com/lbozhilova/bcl2l13-mtdna-selection

## References

1. Elliott HR, Samuels DC, Eden JA, Relton CL, Chinnery PF. Pathogenic mitochondrial DNA mutations are common in the general population. Am J Hum Genet [Internet]. 2008;83(2):254–60. Available from: https://www.ncbi.nlm.nih.gov/pubmed/18674747

2. Stewart JB, Larsson NG. Keeping mtDNA in Shape between Generations. PLoS Genet. 2014;

3. Hagström E, Freyer C, Battersby BJ, Stewart JB, Larsson NG. No recombination of mtDNA after heteroplasmy for 50 generations in the mouse maternal germline. Nucleic Acids Res. 2014;

4. Burr SP, Pezet M, Chinnery PF. Mitochondrial DNA Heteroplasmy and Purifying Selection in the Mammalian Female Germ Line. Dev Growth Differ. 2018;60(1):21–32.

5. Filograna R, Koolmeister C, Upadhyay M, Pajak A, Clemente P, Wibom R, et al. Modulation of mtDNA copy number ameliorates the pathological consequences of a heteroplasmic mtDNA mutation in the mouse. Sci Adv. 2019;5(4).

6. Zhang H, Esposito M, Pezet MG, Aryaman J, Wei W, Klimm F, et al. Mitochondrial DNA heteroplasmy is modulated during oocyte development propagating mutation transmission. Sci Adv [Internet]. 2021 Dec 10 [cited 2022 Jan 7];7(50):5657. Available from: https://www.science.org/doi/abs/10.1126/sciadv.abi5657

7. Kauppila JHK, Baines HL, Bratic A, Simard ML, Freyer C, Mourier A, et al. A Phenotype-Driven Approach to Generate Mouse Models with Pathogenic mtDNA Mutations Causing Mitochondrial Disease. Cell Rep [Internet]. 2016 Sep 13 [cited 2021 Oct 19];16(11):2980–90. Available from: http://www.cell.com/article/S2211124716311019/fulltext

8. Stewart JB, Freyer C, Elson JL, Larsson NG. Purifying selection of mtDNA and its implications for understanding evolution and mitochondrial disease. Nat Rev Genet 2008 99 [Internet]. 2008 Sep [cited 2022 Jan 8];9(9):657–62. Available from: https://www.nature.com/articles/nrg2396

9. Altshuler-Keylin S, Kajimura S. Mitochondrial homeostasis in adipose tissue remodeling. Sci Signal [Internet]. 2017 Feb 28 [cited 2021 Oct 19];10(468). Available from: https://www.science.org/doi/abs/10.1126/scisignal.aai9248

10. Villa E, Marchetti S, Ricci J-EE. No Parkin Zone: Mitophagy without Parkin. Trends Cell Biol [Internet]. 2018 Nov 1 [cited 2021 Oct 19];28(11):882–95. Available from: http://www.cell.com/article/S0962892418301259/fulltext

11. Ma X, McKeen T, Zhang J, Ding W-X. Role and Mechanisms of Mitophagy in Liver Diseases. Cells 2020, Vol 9, Page 837 [Internet]. 2020 Mar 31 [cited 2021 Oct 19];9(4):837. Available from: https://www.mdpi.com/2073-4409/9/4/837/htm

12. Narendra D, Tanaka A, Suen D-F, Youle RJ. Parkin is recruited selectively to impaired mitochondria and promotes their autophagy. J Cell Biol. 2008 Dec 1;183(5):795–803.

13. Pickrell AM, Huang C-H, Kennedy SR, Ordureau A, Sideris DP, Hoekstra JG, et al. Endogenous Parkin Preserves Dopaminergic Substantia Nigral Neurons following Mitochondrial DNA Mutagenic Stress. Neuron [Internet]. 2015 Jul 15 [cited 2021 Oct 19];87(2):371–81. Available from: http://www.cell.com/article/S0896627315005978/fulltext

14. Murakawa T, Yamaguchi O, Hashimoto A, Hikoso S, Takeda T, Oka T, et al. Bcl-2-like protein 13 is a mammalian Atg32 homologue that mediates mitophagy and mitochondrial fragmentation. Nat Commun 2015 61 [Internet]. 2015 Jul 6 [cited 2021 Oct 19];6(1):1–14. Available from: https://www.nature.com/articles/ncomms8527

15. Okamoto K, Kondo-Okamoto N, Ohsumi Y. Mitochondria-Anchored Receptor Atg32 Mediates Degradation of Mitochondria via Selective Autophagy. Dev Cell. 2009 Jul 21;17(1):87–97.

16. Kanki T, Wang K, Cao Y, Baba M, Klionsky DJ. Atg32 Is a Mitochondrial Protein that Confers Selectivity during Mitophagy. Dev Cell. 2009 Jul 21;17(1):98–109.

17. Yan J, Kuroyanagi H, Kuroiwa A, Matsuda YI, Tokumitsu H, Tomoda T, et al. Identification of Mouse ULK1, a Novel Protein Kinase Structurally Related toC. elegansUNC-51. Biochem Biophys Res Commun. 1998 May 8;246(1):222–7.

18. Lane JD, Korolchuk VI, Murray JT, Zachari M, Ganley IG. The mammalian ULK1 complex and autophagy initiation. Essays Biochem [Internet]. 2017 Dec 12 [cited 2021 Oct 19];61(6):585–96. Available from: /essaysbiochem/article/61/6/585/78474/The-mammalian-ULK1-complex-and-autophagy

19. Chang C, Jensen LE, Hurley JH. Autophagosome biogenesis comes out of the black box. Nat Cell Biol 2021 235 [Internet]. 2021 Apr 26 [cited 2021 Oct 19];23(5):450–6. Available from: https://www.nature.com/articles/s41556-021-00669-y

20. Lieber T, Jeedigunta SP, Palozzi JM, Lehmann R, Hurd TR. Mitochondrial fragmentation drives selective removal of deleterious mtDNA in the germline. Nat 2019 5707761 [Internet]. 2019 May 15 [cited 2021 Oct 17];570(7761):380– 4. Available from: https://www.nature.com/articles/s41586-019-1213-4

21. Yan J, Kuroyanagi H, Tomemori T, Okazaki N, Asato K, Matsuda Y, et al. Mouse ULK2, a novel member of the UNC-51-like protein kinases: unique features of functional domains. Oncogene 1999 1843 [Internet]. 1999 Oct 21 [cited 2021 Oct 19];18(43):5850–9. Available from: https://www.nature.com/articles/1202988

22. GTEx Portal [Internet]. [cited 2021 Oct 19]. Available from: https://gtexportal.org/home/

23. The Human Protein Atlas [Internet]. [cited 2021 Oct 19]. Available from: https://www.proteinatlas.org/

24. Ma B, Lee T-L, Hu B, Li J, Li X, Zhao X, et al. Molecular characteristics of early-stage female germ cells revealed by RNA sequencing of low-input cells and analysis of genome-wide DNA methylation. DNA Res [Internet]. 2019 Apr 1 [cited 2021 Oct 19];26(2):105–17. Available from: https://academic.oup.com/dnaresearch/article/26/2/105/5263760

25. Burgstaller JP, Johnston IG, Jones NS, Albrechtová J, Kolbe T, Vogl C, et al. MtDNA Segregation in Heteroplasmic Tissues Is Common InVivo and Modulated by Haplotype Differences and Developmental Stage. Cell Rep [Internet]. 2014 Jun 26 [cited 2022 Apr 30];7(6):2031–41. Available from: http://www.cell.com/article/S2211124714003957/fulltext

26. Wei W, Tuna S, Keogh MJ, Smith KR, Aitman TJ, Beales PL, et al. Germline selection shapes human mitochondrial DNA diversity. Science (80-) [Internet]. 2019 May 24 [cited 2022 Aug 23];364(6442). Available from: https://www.science.org/doi/10.1126/science.aau6520

27. Wallace DC. Mitochondrial DNA mutations in disease and aging. Environ Mol Mutagen. 2010 Jun;51(5):440–50.

28. Ross JM, Stewart JB, Hagström E, Brené S, Mourier A, Coppotelli G, et al. Germline mitochondrial DNA mutations aggravate ageing and can impair brain development. Nat 2013 5017467 [Internet]. 2013 Aug 21 [cited 2022 Aug 1];501(7467):412–5. Available from: https://www.nature.com/articles/nature12474

29. Keogh M, Chinnery PF. Hereditary mtDNA heteroplasmy: A baseline for aging? Cell Metab [Internet]. 2013 Oct 1 [cited 2022 Aug 23];18(4):463–4. Available from: http://www.cell.com/article/S1550413113003847/fulltext

30. Cheong H, Lindsten T, Wu J, Lu C, Thompson CB. Ammonia-induced autophagy is independent of ULK1/ULK2 kinases. Proc Natl Acad Sci U S A [Internet]. 2011 Jul 5 [cited 2022 May 1];108(27):11121–6. Available from: http://www.pnas.org/cgi/doi/10.1073/pnas.1107969108

31. Jeedigunta SP, Minenkova A V., Palozzi JM, Hurd TR. Avoiding Extinction: Recent Advances in Understanding Mechanisms of Mitochondrial DNA Purifying Selection in the Germline. https://doi.org/101146/annurev-genom-121420-081805 [Internet]. 2021 Aug 31 [cited 2022 Sep 1];22:55–80. Available from: https://www.annualreviews.org/doi/abs/10.1146/annurev-genom-121420-081805

32. Muller HJ. The relation of recombination to mutational advance. Mutat Res Mol Mech Mutagen. 1964 May 1;1(1):2–9.

33. Von Coelln R, Thomas B, Savitt JM, Lim KL, Sasaki M, Hess EJ, et al. Loss of locus coeruleus neurons and reduced startle in parkin null mice. Proc Natl Acad Sci U S A [Internet]. 2004 Jul 20 [cited 2022 Jul 21];101(29):10744–9. Available from: https://www.pnas.org/doi/abs/10.1073/pnas.0401297101

34. Lewandoski M, Meyers EN, Martin GR. Analysis of Fgf8 gene function in vertebrate development. Cold Spring Harb Symp Quant Biol. 1997;62:159–68.

